# Tracing Siderophore Precursors to Primary Metabolism for Ecological Applications

**DOI:** 10.1101/2025.04.30.651379

**Authors:** Ruolin He, Jiazheng Xu, Jiqi Shao, Yanzhao Wu, Shizheng Tian, Zihao Yang, Xuejian Li, Haoran Chen, Long Qian, Zhong Wei, Shaohua Gu, Zhiyuan Li

## Abstract

Microbial fitness depends on balancing between primary metabolism supporting basic survival, and secondary metabolism producing metabolites for environmental adaptation. These two systems are interconnected, with most secondary metabolites deriving from primary metabolic precursors. However, systematic frameworks for studying their relationships across diverse microbes remain limited. Using siderophores—iron-chelating secondary metabolites crucial for microbial competition—as a model, we developed a stepwise “siderophore–monomer–precursor–pathway analysis” framework to trace the biosynthetic connections from siderophore structures to primary metabolism. We applied this framework to an expanded SIDERITE database containing 1018 siderophore structures. To demonstrate practical applications, we identified specific precursors utilized by the beneficial rhizobacterium *Bacillus amyloliquefaciens* but not by the plant pathogen *Ralstonia solanacearum*. Supplementing these precursors under iron-limited conditions enhanced both siderophore production in *B. amyloliquefaciens* and its inhibitory activity against *R. solanacearum*. This work establishes a direct link between primary metabolism and siderophore-mediated microbial interactions, offering new strategies for pathogen suppression through targeted metabolic interventions.

## Introduction

Microbial fitness depends on balancing efficient metabolism with environmental adaptation. This balance manifests in the relationship between primary metabolism—supporting growth, replication, and energy generation—and secondary metabolism, which produces specialized compounds for defense, competition, and communication such as antibiotics, siderophores, and signaling molecules^1^. These two metabolic systems are fundamentally interconnected: most secondary metabolites derive from primary metabolic precursors, including amino acids, acetyl-CoA, and the tricarboxylic acid (TCA) cycle intermediates^2^. Acetyl-CoA flows into polyketide and terpene biosynthesis, while amino acids serve as essential substrates for non-ribosomal peptides (NRPs) and ribosomally synthesized and post-translationally modified peptides (RiPPs)^2^. This metabolic interconnection has become central to metabolic engineering strategies. Researchers have enhanced antibiotic production by increasing amino acid precursor supply for NRP-derived compounds like bacitracin and daptomycin^3,4^. Similarly, redirecting carbon flux through suppression of competing pathways has substantially increased secondary metabolite yields^5,6^. However, these approaches typically remain species- and pathway-specific. Despite the fundamental importance of precursor flow, systematic frameworks for studying primary-secondary metabolism connections across diverse systems remain underdeveloped. A broadly applicable, ecologically relevant model is needed to explore these relationships more comprehensively.

Siderophores provide an ideal model for examining the connection between primary and secondary metabolism in microbes. Iron is essential for nearly all organisms, serving as a cofactor in redox enzymes, a structural component in cytochromes, and a building block for iron-sulfur clusters involved in energy metabolism^7^. However, under aerobic, neutral pH conditions, iron becomes insoluble and largely unavailable to cells^8^. This scarcity drives intense microbial competition, particularly in complex environments like the rhizosphere, where microbes depend on siderophores to secure this limiting resource^9^. Siderophores are specialized secondary metabolites that chelate Fe³⁺ from the environment, facilitating uptake through membrane receptors^10^. Their ecological importance is matched by their structural diversity: most siderophores are synthesized by NRPS, one of the most versatile biosynthetic pathways responsible for assembling a wide range of secondary metabolites^11^. Others are produced via NRPS-independent siderophore (NIS) pathways that utilize small-molecule intermediates and alternative biosynthetic mechanisms^12^. This diversity enhances functional adaptation but complicates efforts to trace and compare siderophore biosynthesis across organisms. A systematic approach is therefore needed to decipher the siderophore biosynthesis and connect them to primary metabolism.

To support systematic exploration of siderophore biosynthesis, we previously developed SIDERITE, the most comprehensive curated database of siderophore structures to date^13^. In its 2024 release, SIDERITE contained 707 unique siderophores, each annotated with structural information (including SMILES), functional groups, producing organisms, molecular formula and weight, and classifications based on structural similarity. Unlike general natural product databases, SIDERITE focuses specifically on siderophores and undergoes regular updates, and is the currently only available siderophore database. However, this initial version of SIDERITE treated primary metabolism as a separate domain, without incorporating the biochemical connections between siderophores and the primary metabolic pathways that supply their building blocks. Meanwhile, the modular architecture of many siderophores—particularly those assembled by NRPS—makes it possible to deconstruct their biosynthetic logic. With appropriate analytical tools, the well-structured digital repository in SIDERITE enables quantitative surveys that trace each siderophore’s monomeric components back to their metabolic origins, effectively bridging secondary and primary metabolism through structure-based inference.

In this study, we updated the SIDERITE database in December 2025, expanding the collection from 707 to 1018 unique siderophore structures. Building on this enhanced dataset, we developed a systematic framework to trace the connection between siderophores and primary metabolism through a stepwise siderophore–monomer–precursor–pathway analysis. We used the rBAN tool to deconstruct digitalized siderophore structures into their monomeric components, identified the corresponding biosynthetic precursors through literature mining and BioNavi-NP predictions, and mapped these to known metabolic pathways using KEGG and MetaCyc. This approach provides a quantitative route to explore how secondary metabolites are rooted in primary metabolic networks across diverse microbial systems. To demonstrate the practical utility of this framework, we tested whether supplying specific precursors could enhance beneficial microbial activity in the rhizosphere. We identified siderophore precursors that are utilized by the probiotic bacterium *Bacillus amyloliquefaciens* but not by the soil-borne pathogen *Ralstonia solanacearum*. Supplementing these precursors under iron-limited conditions increased siderophore production in *B. amyloliquefaciens* and enhanced its inhibitory effect on *R. solanacearum*. Together, our findings establish a direct, functional link between primary metabolism and siderophore-driven microbial interactions, offering new strategies for shaping microbial communities and suppressing plant pathogens through targeted metabolic interventions.

## Result

### Update of the SIDERITE database

SIDERITE is a comprehensive siderophore structure database^13^ that we update at least annually since its initial 2023 release, with the most recent update in December 2025. The latest version added 834 new siderophore records with 315 unique structures compared to the May 2024 version (Fig. 1 and Table S1-S2). These additions were collected through a combination of large language model and manual curation^14^. The update includes 95 new monomer, 135 new species, and 46 genera (Fig. 1). Especially, it now covers a new fungal phylum Zoopagomycota (Rhizoferrin in *Basidiobolus microsporus*). Our team’s direct collection efforts now represent the primary source of siderophore records, accounting for 63.41% (1128/1779) of the database. The number of structurally-similar siderophore clusters has increased from 26 to 39, though Clusters 1-4 remain dominant, representing 91.75% of all siderophore structures. In addition, 4 structures, 10 monomers, 12 species, and 4 genera were removed in the latest version due to structural correction. Additional details are available in the update log ( https://github.com/RuolinHe/SIDERITE/issues/1).

**Fig. 1:**
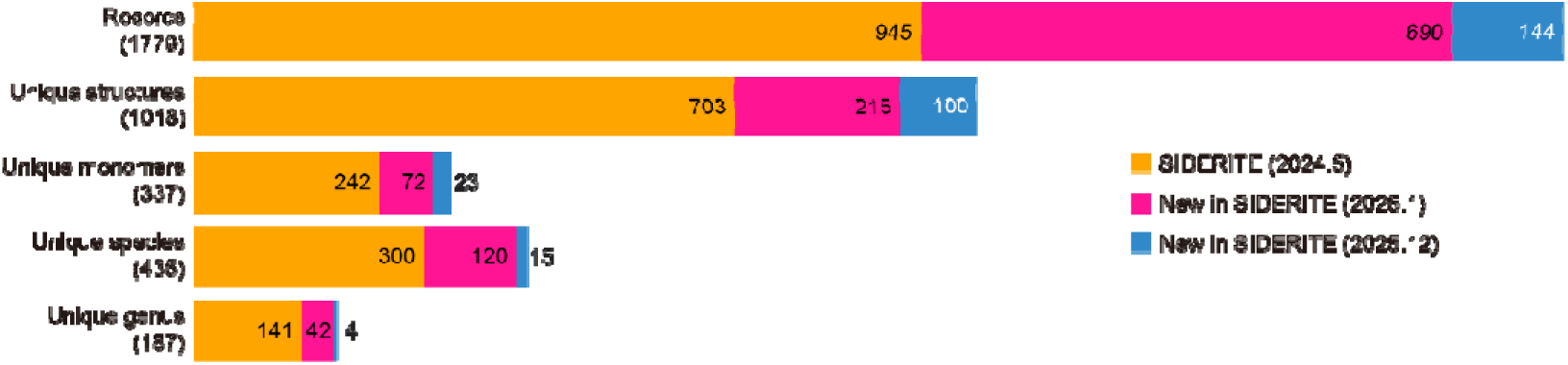
Quantitative overview of updates to SIDERITE in comparison with the previous version. The horizontal bar chart illustrates the growth of the SIDERITE database between the May 2024 version (2024.5, orange), January 2025 update (2025.1, pink) and December 2025 update (2025.12 blue). The database expanded across multiple dimensions: total records increased from 945 to 1,779 (adding 834 new records); unique structures grew from 703 to 1018 (315 new structures); unique monomers increased from 242 to 337 (95 new additions); unique species expanded from 300 to 435 (135 new species); and unique genera grew from 141 to 187 (46 new genera).

Digitalized records enable temporal statistics of siderophore discovery (Fig. S1). Research in this field has accelerated significantly, with approximately 30-40 new siderophores discovered annually since 2010. Our analysis revealed that certain siderophore definitions have evolved over time, with SIDERITE providing comprehensive, up-to-date information valuable to researchers who may not track all developments in this rapidly advancing field (Fig. S2-S3). Despite the substantial database expansion, the properties of siderophores remain largely unchanged after the update (Fig. S4). Bacteria continue to be the primary known siderophore producers, accounting for 88.16% of unique structures. The main biosynthetic pathways remain NRPS (55.50%) and NIS (23.87%). The hybrid NIS/PKS pathway is newly verified by literature, with Amodesmycins being the only reported example^15^. The “Other” type includes siderophore not synthesized by NRPS, NIS, PKS, or their hybrid, such as 2,3-dihydroxybenzoic acid and salicylic acid, plant siderophores derived from nicotianamine, and siderophores with 2-nitrosophenol ligands.

All details of update are recorded in our Google Group (https://groups.google.com/g/siderite-database/c/lboRnKWnxZs) and GitHub (https://github.com/RuolinHe/SIDERITE/issues/1). The history of siderophore discovery, property statistics, and classification of siderophores are simultaneously updated on GitHub Wiki (https://github.com/RuolinHe/SIDERITE/wiki). As SIDERITE is a natural product database, it doesn’t include artificial siderophores, those synthesized by supplementing precursors, or variants produced by gene knock-out strains, though we record their names and references as supplements (Table S3). We also list metabolites containing typical ligands that haven’t been reported as siderophores (mostly antibiotics) as potential siderophores in our supplementary materials. Users can download the latest database and supplementary files from Zenodo (https://doi.org/10.5281/zenodo.18327880).

### Distribution of siderophore monomers

The biosynthesis of siderophores is a major focus in this field. Siderophores form by ordered assembly of related substrates. NRPS pathways primarily utilize amino acids as substrates, while NIS pathway prefers carboxylic acids^12^. Additionally, different species often exhibit different substrate preferences^16^. Creating a comprehensive statistical overview of these substrate preferences helps to predict the biosynthetic enzyme and producers for specific siderophores.

To analyze substrate preferences in siderophores, we used retro-biosynthetic analysis of non-ribosomal peptides (rBAN)^17^. rBAN splits common bonds in siderophore synthesized by NRPS or NIS pathways, like peptide bonds and ester bonds, to identify their component monomers. These monomers represent either the original substrates or their derivatives modified during biosynthesis. Since accurate substrate information requires detailed experiments on the substrate specificity of each module in the biosynthetic gene clusters (BGC), we use monomers as approximations of the substrates.

Our analysis successfully resolved 92.55% of monomers across all siderophores with our customized monomer database (Table S4-S5). The resolution rate was particularly high for NRPS siderophores (95.47% across 565 structures) and NIS siderophores (95.47% across 244 structures). PKS and hybrid-type siderophores generally showed lower resolution rates: 32.36% for 11 PKS siderophores, 83.67% for 150 hybrid NRPS/PKS siderophores, 82.67% for 20 hybrid NRPS/NIS siderophores, and 82.50% for 2 hybrid NIS/PKS siderophores. The “other type” category of siderophores had a resolution rate of 87.13% across 27 structures. Overall, given that NRPS and NIS pathways account for the majority of siderophore biosynthesis, their high resolution rates provide sufficient representation for our comprehensive substrate analysis.

Among the top 30 monomers in siderophores, most function either as scaffolds or chelating components (Fig. S5 and Fig. 2). The most common monomer, serine (Ser), forms the backbone scaffold in catechol siderophores such as enterobactin, turnerbactin, chrysobactin, and vanchrobactin. Similarly, succinate (Suc), the second most common monomer, provides the structural framework for NIS siderophores like desferrioxamines. From the perspective of iron chelating, Orn derivatives such as Ac-OH-Orn, OH-Orn, Fo-OH-Orn, OH-cOrn, and NAc-OH-Orn provide hydroxamate ligands, diOH-Bz and ChrP provide catechol ligands, and OH-Asp and Citric Acid (CitA) provide alpha-hydroxycarboxylate ligands. Notably, all chelating ligands come from non-proteinogenic amino acids, explaining why ribosomal pathway cannot synthesize siderophores. These findings align with Maude Pupin’s 2010 research on 88 siderophores^18^, though our more comprehensive dataset reveals different priorities among the top monomers.

**Fig. 2:**
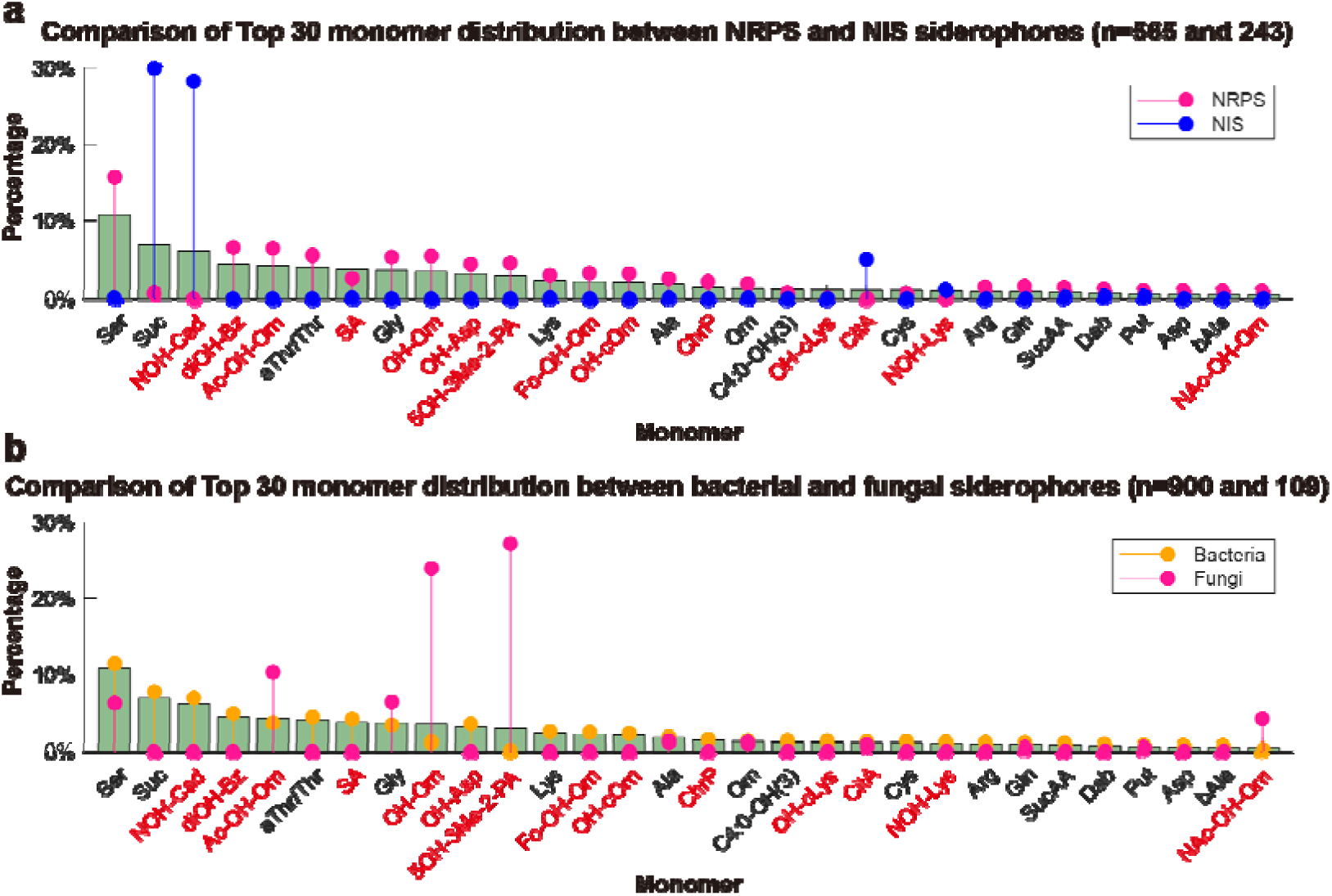
Monomer distribution patterns in siderophores by biosynthetic pathway and producer organism. **a** Comparison of monomer distribution between NRPS (n=565) and NIS (n=243) siderophores. The graph shows the percentage frequency of the top 30 monomers in the database. Abbreviations include standard amino acids (Ser, Gly, Lys, etc.) and specialized monomers like diOH-Bz (2,3-dihydroxybenzoic acid), Ac-OH-Orn (N-acetyl-HydroxyOrnithine), and ChrP (pyoverdin chromophore). See Table S4 for all abbreviations. Monomers that form chelating ligands are highlighted in red. Only the top 30 most frequently occurring monomers are shown. Unknown monomer is hidden. **b** Comparison of monomer distribution between bacterial (n=900) and fungal (n=109) siderophores. The legend is same as **a**.

Siderophores from NRPS and NIS pathways show apparent different monomer compositions (Fig. 2a). NRPS siderophores prefer Ser (scaffold), Orn derivatives (chelation), aThr/Thr (scaffold), diOH-Bz (chelation), Gly (scaffold), and OH-Asp (chelation). In contrast, NIS siderophores prefer Suc (scaffold), NOH-Cad (chelation), and CitA (chelation). The distinct monomer distribution between NRPS and NIS siderophores indicate different substrate pools.

Bacterial and fungal siderophores also exhibit different monomer preferences. Since bacteria produce most siderophores in SIDERITE (88.16%, Fig. S4), bacterial siderophore monomer distribution closely resembles that of the entire database (Fig. 2b and Fig. S5f). Fungal siderophores show a more concentrated pattern, with 5OH-3Me-2-PA and OH-Orn (both derived from substrate N5-anhydromevalonyl-N5-hydroxy-L-ornithine (AMHO) accounting for 51.11% fungal monomers. Overall, fungal siderophores prefer three Orn derivatives (AMHO, Ac-OH-Orn, and NAc-OH-Orn) for chelation, along with Ser and Gly for scaffold (Fig. 2b). These five monomers constitute 73.85% fungal siderophore monomers, suggesting more conserved biosynthetic pathways in fungi compared to bacteria.

Pyoverdine, a characteristic NRPS siderophore primarily produced by *Pseudomonas*, has been extensively studied in the siderophore field^19^. In SIDERITE, the 79 pyoverdine structures constitute the fourth-largest cluster and represent the most significant contribution from a single genus. Due to pyoverdine’s structural diversity, we carefully ensured that it did not bias our comparison between NIS and NRPS siderophores (Fig. 2a), by examining their monomer distribution separately (Fig. S5c-S5d). Similar to other NRPS siderophores, pyoverdines preferentially utilize Ser (18.72%) and Orn derivatives (16.63%, Fig. S5d). Additionally, all pyoverdines have a characteristic chromophore such as ChrP.

### Connecting secondary metabolism with primary metabolism

Siderophore synthesis depends on substrates derived from precursors from primary metabolic pathways, including TCA cycle. This connection between siderophores and primary metabolism links secondary metabolites to essential cellular processes. Tracing siderophore precursors back into primary metabolism provides insights into their relationship, an area of significant scientific interest^20–22^.

By analyzing siderophores at their monomer level, we trace their precursors in primary biosynthetic pathways (Fig. 3a-3c and Table S6-S7). Using established databases such as KEGG^23^ and MetaCyc^24^, we compiled a comprehensive set of precursor molecules, including the 20 proteinogenic amino acids, two non-proteinogenic amino acids (Orn and Dab), and various metabolites involved in primary metabolic pathways such as glycolysis, the pentose phosphate pathway (PPP), the TCA cycle, the glyoxylate cycle, and fatty acid biosynthesis and oxidation. Through literature curation and the BioNavi-NP tool^25^, we systematically traced back each siderophore substrate’s synthesis pathway until reaching an established precursors. In total, we identified 39 precursors that generate 332 resolvable monomers (two irresolvable monomers with unknown precursors are left in Table S6). Notably, all amino acids, and Citric Acid and succinate, function both as precursors and as direct substrates for siderophore synthesis. These 39 precursors span all curated primary metabolism pathways (Fig. 3d), with the TCA cycle and fatty acid biosynthesis/oxidation providing the greatest variety of precursors (Fig. 3d). The PPP primarily contributes His and three aromatic amino acids (Trp, Phe, and Tyr).

**Fig. 3:**
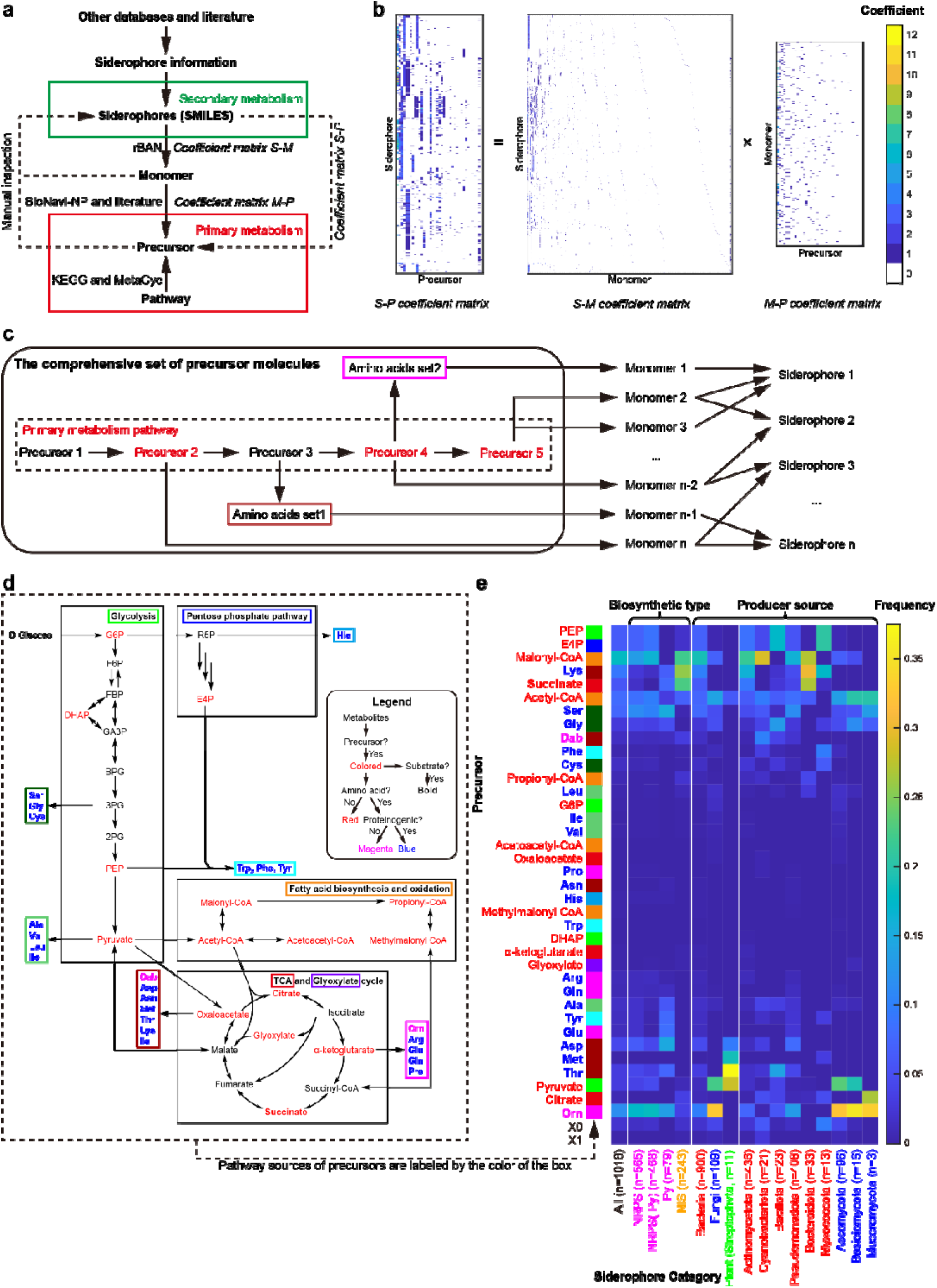
Connecting primary metabolism to siderophore biosynthesis. **a** The workflow of siderophore analysis. After collecting siderophores structures, monomers are identified by rBAN, precursors are determined by BioNavi-NP and the literature, and pathway source are retrieved from KEGG and MetaCyc. Secondary metabolism (green) connects to primary metabolism (red) by coefficient matrices that quantify relationships between siderophores, monomers, and precursors. **b** Calculation of coefficient relationships. The siderophore-precursor relationship (Coefficient matrix S-P) equals the multiplication of siderophore-monomer (Coefficient matrix S-M) and monomer-precursor (Coefficient matrix M-P) matrices. Coefficient values range from 0 to 12 **c** The schematic diagram of precursor tracking. We defined a comprehensive set of precursor molecules including 20 proteinogenic amino acids, 2 non-proteinogenic amino acids (Orn and Dab), and key metabolites from primary pathways. The precursor of each siderophore monomer is identified as the first match from this set during tracking **d** Distribution of siderophore precursors in the primary metabolism. For clarity, only the core parts of the pentose phosphate pathway and fatty acid biosynthesis and oxidation are shown. Precursors are color-coded in their texts by types: non-amino acid precursors (red), proteinogenic amino acids (blue), and non-proteinogenic amino acids (magenta). Bold fonts indicate precursors that also serve as direct substrates for siderophore synthetases. **e** Precursors frequency across different siderophores classifications. Data is shown for biosynthetic types (NRPS in green, NIS in yellow) and producer sources (bacteria in red, fungi in blue, plants in green), with further breakdown by phylum. X0 and X1 represent unknown precursors. Precursors are arranged by hierarchical clustering. Their pathway sources are color-coded with filled boxes around precursors. The colors come from box colors in **d**. Especially for precursors from TCA and glyoxylate cycles, only glyoxylate is colored in purple (glyoxylate cycle), while other precursors are colored in red (TCA cycle). Abbreviations of precursors refer to Table S11.

The contribution of different precursors to siderophore biosynthesis varies substantially (Table S8). To quantify their importance, we defined two metrics: “occurrence frequency,” which indicates the proportion of monomers or siderophores containing a precursor at least once and indicates pervasiveness, and “utilization frequency,” which reflects the total number of times a precursor is used relative to all precursor units and quantifies overall contributes (see Methods for details).

Among monomers, malonyl-CoA exhibits the highest occurrence frequency, serving as a key building block for fatty acid synthesis, followed by acetyl-CoA, which supports even-numbered straight-chain fatty acid biosynthesis and acetylation (Table S9). In siderophores, ornithine (Orn) and serine (Ser) show the highest occurrence frequencies, underscoring their critical roles in siderophore assembly. Notably, precursors with low derivative diversity (≤8 derivatives) but high occurrence frequency in siderophores (>9%) correspond to monomers functioning as structural scaffolds (Table S9, Fig. 2), highlighting their pervasiveness

Precursor utilization frequency reveals distinct preferences in siderophore biosynthesis across different biosynthetic pathways and microbial producers (Table S10 and Fig. 3e). Except malonyl-CoA and acetyl-CoA, NRPS siderophores have their substrates predominantly derived from Orn and Ser, while NIS siderophores substrates favor Lys and Suc precursors. Pyoverdines, a major NRPS siderophore group, frequently utilize Asp, which provides alpha-hydroxycarboxylate ligand in OH-Asp for iron chelation. From the producer perspective, bacterial siderophores prefer Orn and Ser, while fungal siderophores prefer Orn and pyruvate. Plant siderophores, found only in Streptophyta phylum, prefer Thr, pyruvate, Met, and Asp. Producers from different phyla show characteristic precursor usage patterns (Fig 3e, 9-17 columns of right panels). These insights into precursor preferences could inform strategies to inhibit or enhance siderophore production in specific species by targeting particular precursor synthesis pathways

### Targeted Siderophore Precursor Supplementation Enhances Control of a Soil Pathogen

Building on our understanding of precursor preferences in siderophore biosynthesis, we explored practical applications by testing whether targeted precursor supplementation could selectively enhance beneficial microbial activity against pathogens. To facilitate the assay, the strains used should synthesize only one type of siderophore. We selected the beneficial strain *Bacillus amyloliquefaciens* T-5, known for controlling soil-borne bacterial wilt, and the pathogenic *Ralstonia solanacearum* QL-Rs1115 as our experimental system^26,27^. This pair provides an ideal model since each organism produces only one type of siderophore with distinct precursor requirements (Fig. 4a). *B*. *amyloliquefaciens* T-5 produces bacillibactin whose precursors are Thr, Gly, PEP, and E4P. *R*. *solanacearum* QL-Rs1115 produces staphyloferrin B, which utilizes Ser, CitA and Glu as precusors^28^.

**Fig. 4:**
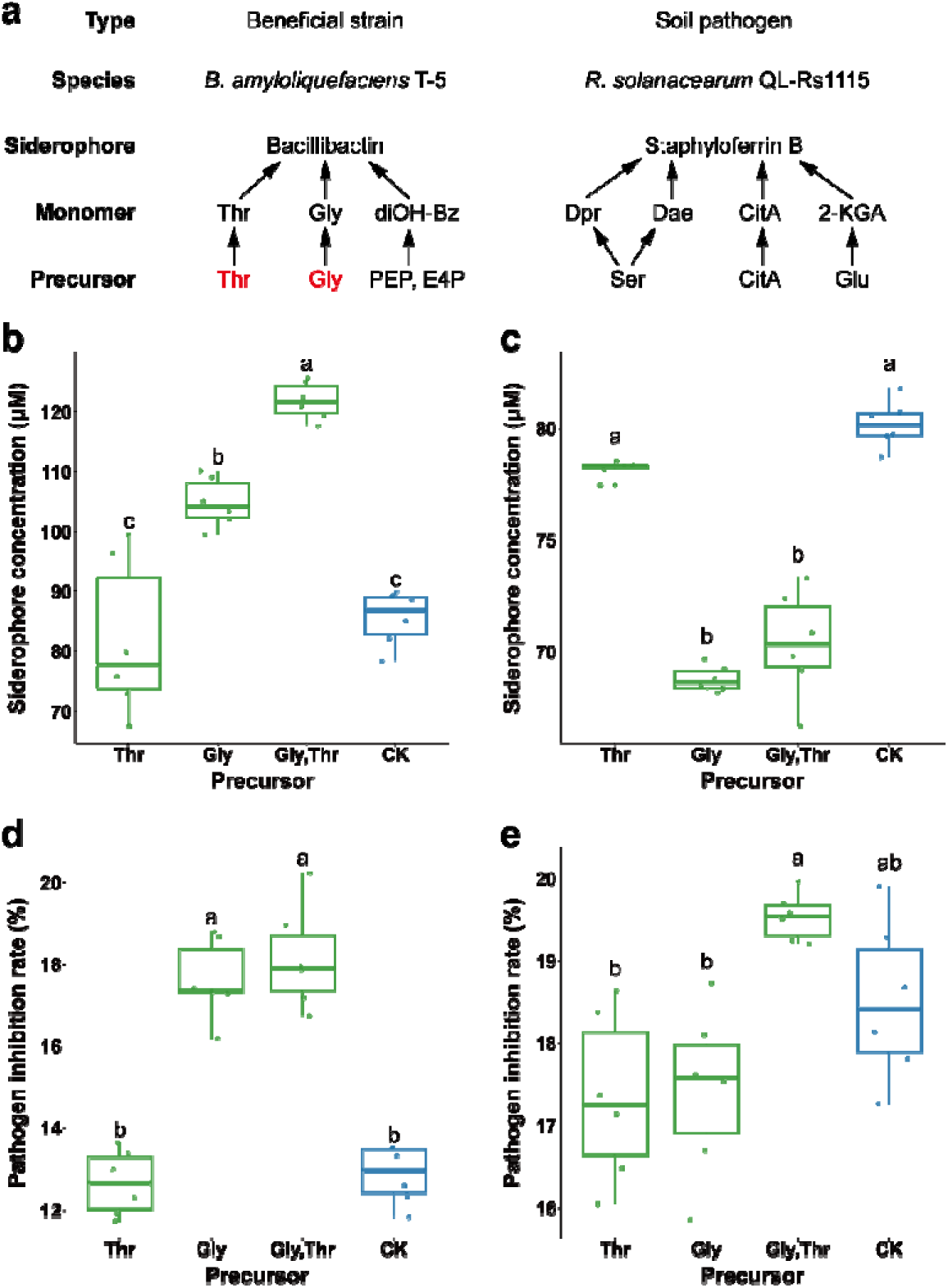
Bacillibactin precursors selectively enhance siderophore production in *Bacillus amyloliquefaciens* and improve its inhibitory activity against *R. solanacearum*. **a** Comparison between two siderophores used in experiments. Supplemented precursors are highlighted in red. **b** Effect of precursors on siderophore production in *B*. *amyloliquefaciens*. **c** Effect of precursors on siderophore production in *R. solanacearum*. **d** Effect of precursors on antagonistic activity of *B*. *amyloliquefaciens* against *R. solanacearum* under iron-limited conditions. **e** Effect of precursors on antagonistic activity of *B*. *amyloliquefaciens* against *R. solanacearum* under iron-rich conditions. Data are presented as box plots where the central line represents the median, the box edges indicate the first and third quartiles, and the whiskers extend to the minimum and maximum values. Different letters indicate statistically significant differences (p<0.05). The x-axis labels “Thr” and “Gly” denote individual supplementation with threonine or glycine, respectively, while “Gly,Thr” indicates co-supplementation (each precursor at a final concentration of 6 mM). “CK” represents the control without precursor addition.

Our experiments revealed that supplementing the growth medium with glycine, a key bacillibactin precursor, significantly increased siderophore production in *B*. *amyloliquefaciens* under the iron-limited condition. The combined addition of both Gly and Thr yielded the highest production increase of approximately 40% (Fig. 4b). In contrast, these same amino acids are not precursors for siderophore biosynthesis in *R. solanacearum*, and their supplementation, either alone or in combination, did not enhance siderophore production in *R. solanacearum* (Fig. 4c). These findings confirm that selective precursor supplementation can realize targeted siderophore-production enhancement.

To assess whether selectively enhanced siderophore production translated to improved pathogen control, we examined bacterial interactions under both iron-limited and iron-rich conditions. Under iron limitation, supplementation alone or combined with threonine significantly enhanced the inhibitory effect of *B. amyloliquefaciens* against *R. solanacearum* (Fig. 4d). However, under iron-rich condition where siderophore production is naturally suppressed, these same supplementations provided no enhancement and sometimes even reduced the inhibitory effect (Fig. 4e). These findings, alongside the siderophore production data in Fig. 4b, demonstrate that strategic supplementation with specific precursors can selectively enhance beneficial microbial activity against pathogens by utilizing the differences in their siderophore biosynthetic pathways.

## Discussion

Siderophores, with their widespread presence, ecological importance, structural diversity, and dependence on primary metabolic precursors, stand out as an ideal model system for exploring the connections between primary and secondary metabolism. In this study, we have taken significant steps to deepen this understanding by enhancing the SIDERITE database, now featuring 1018 unique siderophore structures, and developed a methodology linking their building blocks to primary metabolic pathways. This enhanced framework allowed us to predict whether supplementing specific precursors could selectively boost siderophore production in beneficial bacteria to counteract pathogens. This work makes three key contributions: (1) the first comprehensive database connecting siderophore structures to primary metabolism, (2) a systematic analysis revealing relationships between siderophore synthesis and primary metabolic precursors—addressing a gap left by previous studies focused on individual pathways or species, and (3) a novel “selective precursor supplementation” approach for fine-tuning microbial communities, with potential applications in agricultural disease management. These advances deepen our understanding of microbial metabolism while offering practical applications with real-world impact.

Regulating microbial communities presents both important opportunities and significant challenges. The key to effective regulation lies in exploiting metabolic differences—selective enhancement of beneficial bacteria is only possible when they have distinct metabolic requirements from pathogens. The structural diversity of siderophores provides the foundation for such differentiation, and our “selective precursor supplementation” strategy offers an innovative and safe metabolic intervention for targeted regulation of beneficial microorganisms. By selectively supplementing primary metabolic precursors, we enhance siderophore production in beneficial bacteria, effectively suppressing pathogens. This approach provides significant advantages over genetic engineering by avoiding ecological risks associated with releasing genetically modified strains^29^.

Previous research demonstrates this concept’s viability—glucose and tryptophan supplementation significantly increased anticancer precursor production in yeast^30^, while optimized precursor supply enhanced Avermectin B1a production in Streptomyces^6^. However, successful implementation depends on thorough understanding of the differences in siderophore types and precursor requirements between beneficial microbes and pathogens—information now accessible through our enhanced SIDERITE database.

Our update to the SIDERITE database (May 2024-December 2025) expanded its coverage to 1018 unique siderophore structures. We combined large language models with manual curation to improve collection efficiency, with ChatGPT proving especially valuable in identifying literature lacking typical keywords such as “siderophore” or “novel”. Surprisingly, only 84 of the 315 new structures were reported in 2024-2025, with most coming from previously overlooked historical studies. Though yielding under 10% positive hits, this AI-assisted approach significantly outpaced traditional methods. Moving forward, SIDERITE must evolve from structural descriptions toward a quantitative framework by integrating genomic and metabolomic data. This integration will connect structures to their genetic origins and metabolic pathways, enabling predictive tools for novel siderophore identification and functional characterization. Following this major expansion, SIDERITE will shift to annual updates of 30-40 new structures, supporting both fundamental research and practical applications in biotechnology and related fields.

In future work, building on SIDERITE’s systematic analysis, we can deepen our understanding of the interplay between primary and secondary metabolism, moving beyond the simple concept that primary metabolism merely supplies precursors for secondary metabolites. Our analysis of siderophore monomers reveals that all monomers containing chelating ligands are derived from non-proteinogenic amino acids, while some of the top-ranking monomers are proteinogenic amino acids for scaffolding. This suggests a potential competition or conflict between secondary and primary metabolism for these resources. The extent of this competition likely depends on the proportion of secondary metabolites in the total cellular metabolite pool, which warrants further investigation through quantitative metabolic profiling.

Looking ahead, future siderophore research should focus on advancing our analytical tools, particularly for complex PKS and hybrid siderophore systems where biosynthetic prediction remains challenging. Our work establishes a regulatory framework linking primary metabolic precursors to secondary metabolites and nutritional immunity, creating a foundation for sustainable pathogen control in agriculture. Future studies could investigate the synergistic effects of multiple precursors and optimize regulation strategies based on specific microenvironmental characteristics. Interdisciplinary collaboration—integrating bioinformatics, metabolic engineering, and microbial ecology—will be crucial for translating these discoveries into innovative biotechnological solutions addressing agricultural and medical challenges.

## Method

### Siderophore information acquisition

The updates to the SIDERITE data was updated using a combination of manual collection and ChatGPT assistance^14^. Briefly, we identified papers potentially containing missing siderophore structures, then manually verified their relevance before adding the siderophore information to the database. ChatGPT searched papers published from the 1980s to December 31, 2025, while manual collection extended to January 31, 2025. In total, this update includes papers up to December 31, 2025.

### Annotation of siderophore monomers, precursors and pathways

Monomers of siderophores are split by rBAN^17^. We expanded the original monomer database to cover monomers in siderophores (Table S4). First, we obtained new monomers by “discoveryMode” in rBAN and added new monomers into our customized monomer database after manual verification (Table S5). Some “new” monomers were artifacts because rBAN overly split the bonds (e.g. peptide bonds) in the monomer. For example, N-[2-(2-aminoethylsulfanyl)ethyl]hydroxylamine (NOH-DAT) is overly split into Cysteamine and N-ethylhydroxylamine in Desferrioxamine Te2 and Desferrioxamine Te3. These errors were corrected by a customized Matlab script. The customized monomer database can be found in GitHub (https://github.com/RuolinHe/SIDERITE/tree/main/monomer_analysis).

Precursors of siderophore monomers were sourced from literature or predicted using BioNavi-NP^25^. In total, we identified 39 precursors from 332 resolvable monomers (two irresolvable monomers with unknown precursors are left in Table S6). Precursors’ location in primary biosynthetic pathway (glycolysis, fatty acid biosynthesis and oxidation, pentose phosphate pathway, tricarboxylic acid cycle, and glyoxylate cycle) are identified by KEGG^23^ and MetaCyc^24^ (Table S7).

### Calculation of siderophore-precursor coefficient matrix

After annotation of siderophore monomers and precursors, we obtained two coefficient matrices. For siderophore-monomer (S-M) coefficient matrix, each row is one siderophore and each column is one monomer (Table S5). The coefficient is the number of the specific monomer used in this siderophore. Similarly, for monomer-precursor (M-P) coefficient matrix, each row is one monomer and each column is one precursor (Table S6). The coefficient is the number of the specific precursor used in this monomer. The siderophore-precursor (S-P) coefficient matrix is equal to S-M coefficient matrix multiplied by M-P coefficient matrix (Table S8).

### Occurrence frequency and utilization frequency of precursors

We define two key metrics to quantify the importance of siderophore precursors: **occurrence frequency and utilization frequency.**

- **Occurrence Frequency**: This metric represents the proportion of monomers (or siderophores) in which a specific precursor appears at least once. To calculate it, we first determine the *occurrence count*, which is the number of unique monomers (or siderophores) that contain at least one instance of the precursor. The occurrence frequency is then computed by dividing this occurrence count by the total number of monomers (or siderophores) in the dataset. This measure highlights how widespread a precursor is across different compounds, regardless of how many times it appears within each one.
- **Utilization Frequency**: This metric reflects the proportion of all precursor units across all monomers (or siderophores) that belong to a particular precursor type. We calculate the *utilization count* as the total number of times the precursor appears across all monomers (or siderophores), accounting for multiple instances within the same compound. The utilization frequency is then determined by dividing this utilization count by the sum of the utilization counts for all precursors in the dataset. This measure captures the precursor’s overall quantitative contribution to the composition of the compounds.

Together, these metrics provide complementary perspectives: occurrence frequency indicates the distribution of a precursor across distinct compounds, while utilization frequency quantifies its total usage relative to other precursors.

### Effect of precursors on bacterial siderophore production

The rhizosphere-beneficial bacterium *Bacillus amyloliquefaciens* T-5 (China General Microbiology Culture Collection Center (CGMCC) accession No. 8547), previously isolated from a healthy tomato plant in a wilt diseased field, were streaked onto TSA solid media^26^. Pathogenic bacterium *Ralstonia solanacearum* QL-Rs1115 (GenBank accession GU390462) tagged with the pYC12-mCherry plasmid, isolated from the roots of a wilted tomato plant were streaked onto NA solid media^27^.

Single colonies were subsequently cultured at 30°C. T-5 colonies were inoculated into TSB liquid medium, while *R. solanacearum* colonies were inoculated into NB liquid medium, using 24-well plates with 1.5 mL of medium per well. Cultures were incubated at 30°C and 170 rpm for 16 hours. The bacterial suspensions were then adjusted to an OD_600_ of 0.5 using sterilized saline (0.9 g NaCl dissolved in 100 mL deionized water). Next, 2 μL of T-5 and *R. solanacearum* suspensions were separately inoculated into a 96-well plate containing 178 μL of iron-limited MKB medium. As treatment conditions, 20 μL of precursor solution (Gly, Thr, or Gly+Thr, each at a final concentration of 6 mM) or 20 μL of sterile water (control) were added. Each treatment was conducted in six replicates, and cultures were incubated in a constant-temperature shaker at 30°C and 170 rpm.

After 48 hours incubation, OD_600_ values were measured using a microplate reader (Tecan Infinite Eplex). The bacterial cultures were then centrifuged using a well-plate centrifuge and a 96-well plate filter (3300 rpm, 10 min) to obtain the supernatant. To quantify siderophore production, the sterile supernatant and the sterile MKB medium were respectively mixed with the CAS detection solution at a ratio of 1:1 and left to react at room temperature for 2 hours. OD_630_ values of the sterile supernatant (A) and the sterile MKB medium control (Ar) were measured using a microplate reader. The siderophore relative content (SU) was calculated using the formula:

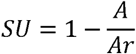

Since both T-5 and *R. solanacearum* produce high levels of siderophores, the bacterial supernatant was diluted to ensure SU < 0.73. The siderophore equivalent (SE) was then calculated using the formula^31^:

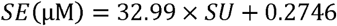

Multiplying by the dilution factor provided a more accurate siderophore concentration.

### Effect of precursors on the inhibitory activity of rhizosphere-beneficial bacteria T-5 against pathogen *R. solanacearum*

2 μL of rhizosphere-beneficial bacteria T-5 and 2 μL of pathogen *R. solanacearum* bacterial suspensions (both adjusted to an OD_600_ of 0.5) were inoculated into 96-well plates containing 178 μL of either iron-limited MKB medium or iron-rich MKB medium (supplemented with FeCl_3_ at a final concentration of 50 μM)^27^. As treatment conditions, 20 μL of precursor solution (Gly, Thr, or Gly+Thr, each at a final concentration of 6 mM) or 20 μL of sterile water (control) were added. A positive control was included by replacing the T-5 bacterial suspension with sterile water. Each treatment was conducted in six replicates. The well plates were incubated in a constant-temperature shaker at 30°C and 170 rpm for 24 hours. Following incubation, OD_600_ values (OD) and red fluorescence intensities (RFP) *R. solanacearum* (excitation wavelength: 587 nm, emission wavelength: 610 nm) were measured using a microplate reader. The inhibitory effect of T-5 and the precursor treatments on pathogen *R. solanacearum* was assessed by calculating the relative biomass of pathogen *R. solanacearum* under co-culture conditions.

The abundance of pathogen *R. solanacearum* was calculated as:

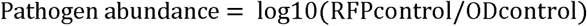

The inhibition rate of pathogen *R. solanacearum* was determined using the formula:

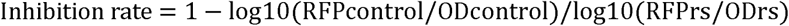

where RFPrs represents the fluorescence intensity of pathogen *R. solanacearum* in the positive control (cultured alone), and RFPcontrol represents the fluorescence intensity of pathogen *R. solanacearum* after treatment.

### Analysis of experiment result

Experimental data were processed using statistical software, including Excel 2016 and the R programming language. Data analysis was performed using ANOVA, and significance was assessed using Tukey’s HSD test.

## Acknowledgements

This work was supported by the National Key R&D Program of China (No. 2024YFA0919500, No. 2020YFA0906901), National Natural Science Foundation of China (No. T2321001). LZ was supported in part by the Peking-Tsinghua Center for Life Sciences.

