## Supplemental information for "Tracing Siderophore Precursors to Primary Metabolism for Ecological Applications"

### The legend of Supplementary Materials

**Supplementary Materials** contains **Figure S1-S5**. See attached Excel files for **Table S1-S9**.

**Figure:**

**Figure S1.** The history of siderophore discovery.

**Figure S2.** The change in the meaning of corynebactin.

**Figure S3.** The change in the meaning of heterobactin A.

**Figure S4.** The statistics of siderophore property.

**Figure S5.** The top 30 monomer distribution of siderophores.

**Table:**

**Table S1.** Detailed information of 1779 siderophore records.

**Table S2.** Detailed information of 1018 unique siderophore structures.

**Table S3.** Non-natural siderophore records.

**Table S4.** The customized monomer database.

**Table S5.** Siderophore-Monomer coefficient matrix.

**Table S6.** Monomer-Precursor coefficient matrix.

**Table S7.** Table of siderophore precursor-pathway pairs.

**Table S8.** Siderophore-Precursor coefficient matrix.

**Table S9.** The utilization frequency of precursors in the monomers and siderophores with different siderophore categories.

**Table S10.** The occurrence frequency of precursors in the monomers and siderophores.

**Table S11.** Abbreviations of siderophore precursors.

**Reference**

### Figure S1

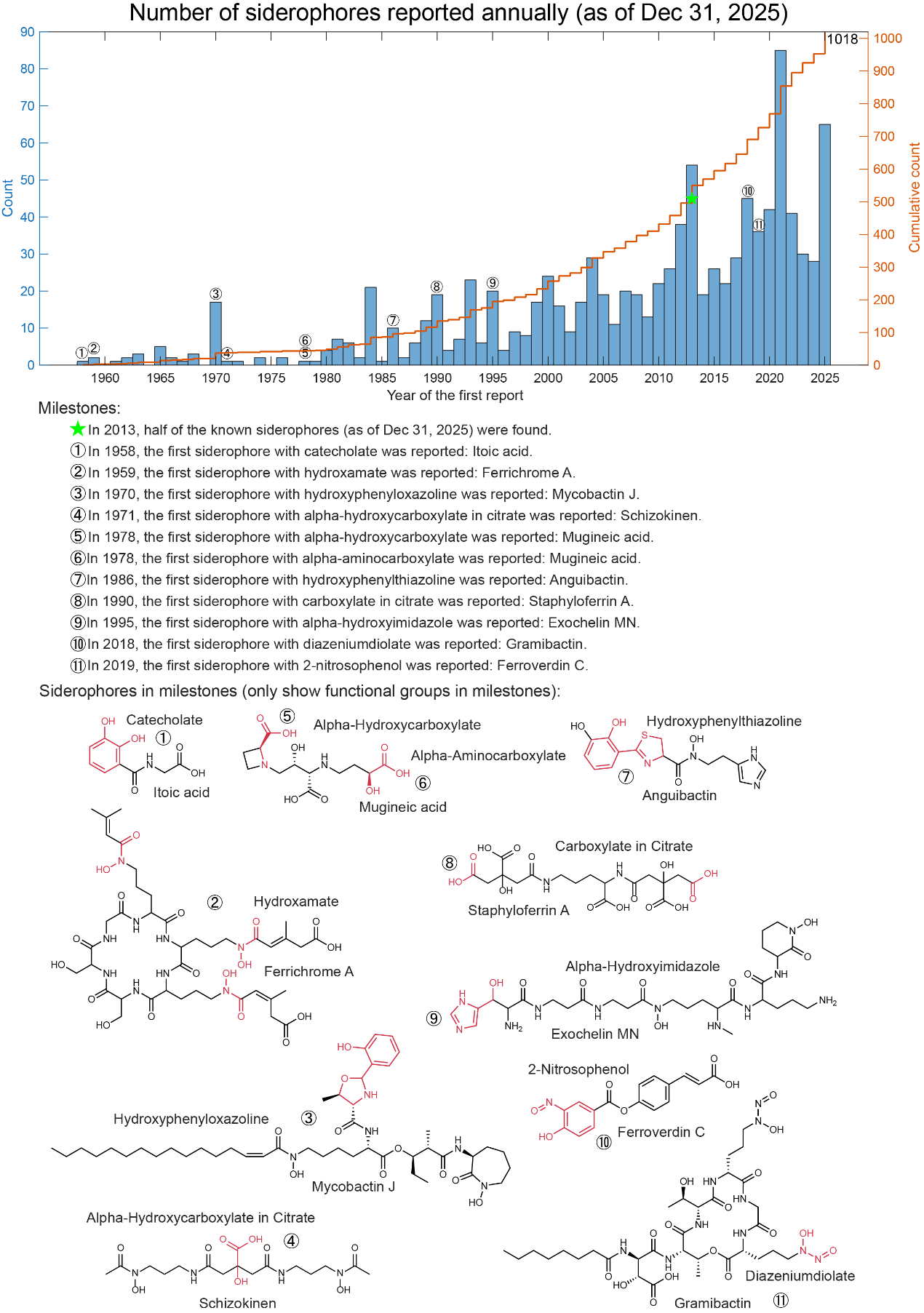

**Figure S1. The history of siderophore discovery.**

In this study, we use the year when the siderophore was first published as the discovery year of this siderophore. The first siderophore is itoic acid produced by *Bacillus subilis* in 1958 (%cite: "Products of “low-iron fermentation” with Bacillus subilis: isolation, characterization and synthesis of 2, 3-dihydroxybenzoylglycine1, 2.). Since that, the number of new siderophores has increased year by year. After 2010, around 30~40 siderophores were discovered every year. However, only 2 of the 11 ligand types were recently discovered, and 9 were discovered before 1995, indicating that the major siderophore ligand types are known. We marked the milestones of siderophore discovery that including the time point when half of all siderophores were discovered, and the time point when 11 main siderophore ligands were first identified. The structures of siderophores mentioned in the milestones are shown in the lower panel. The functional groups (ligands) are highlighted in red.

### Figure S2

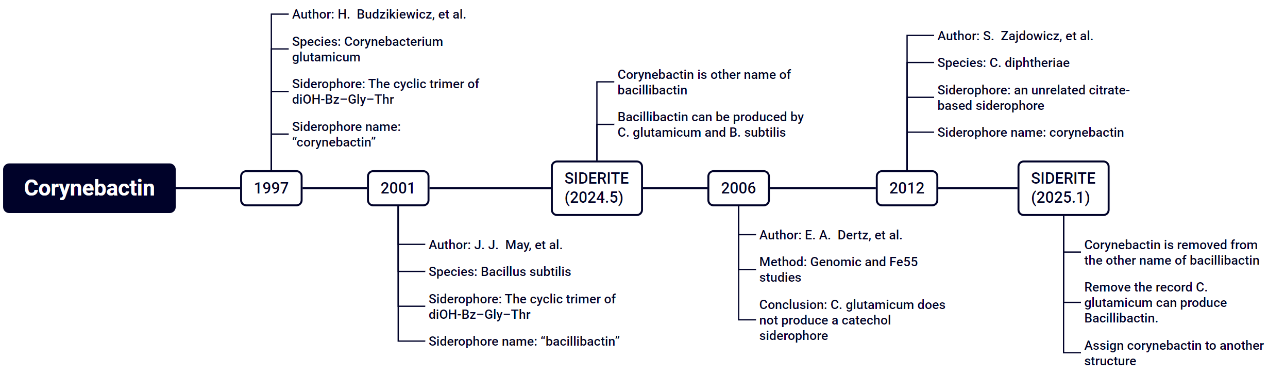

**Figure S2. The change in the meaning of corynebactin.**

First case is about bacillibactin and corynebactin which is mentioned in Zachary L Reitz’s work^1^ (Fig. S). The cyclic trimer of diOH-Bz–Gly–Thr was first reported as “corynebactin” in *Corynebacterium glutamicum* in 1997^2^. And in 2001, the cyclic trimer of diOH-Bz–Gly–Thr was reported in *Bacillus subtilis* and named as “bacillibactin”^3^. In the last version of SIDERITE, bacillibactin and corynebactin are synonym. However, researcher found *C. glutamicum* didn’t produce a catechol siderophore by genomic and Fe^55^ studies in 2006^4^. And “corynebactin” was assigned to another siderophore produced by *C. diphtheriae* in 2012^5^. Therefore, in the latest SIDERITE, corynebactin is removed from the other name of bacillibactin. Then, we removed the record *C. glutamicum* can produce bacillibactin and assigned corynebactin to another structure.

### Figure S3

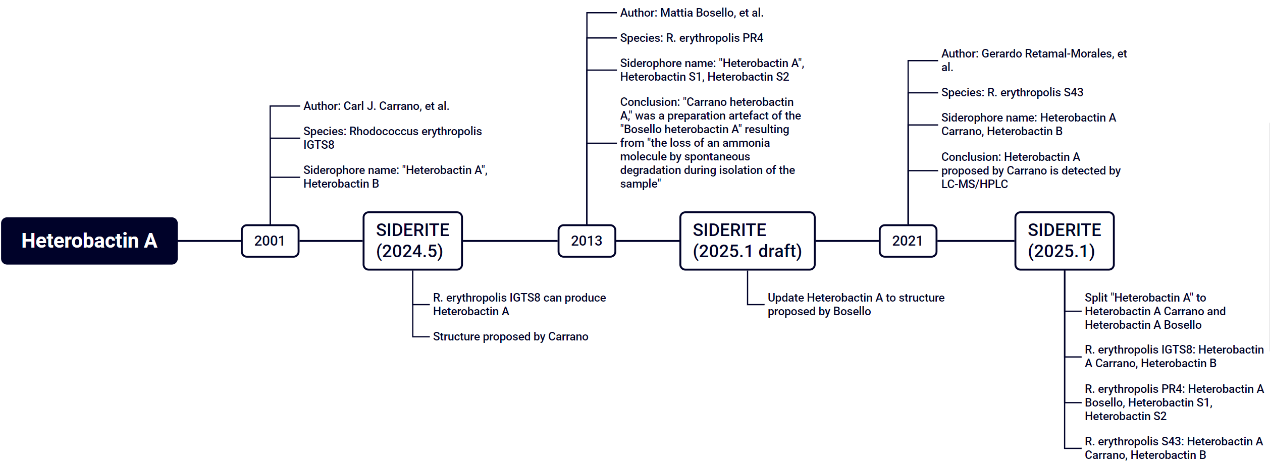

**Figure S3. The change in the meaning of heterobactin A.**

Second case is about heterobactin A (Fig. S). In 2001, “heterobactin A” was first discovered with heterobactin B in *Rhodococcus erythropolis* IGTS8 by Carl J. Carrano, et al^6^. In the last version of SIDERITE, *R. erythropolis* IGTS8 can produce heterobactin A. However, in 2013, Mattia Bosello, et al. reported another “heterobactin A” structure with heterobactin S1 and heterobactin S2 in *R. erythropolis* PR4^7^. Mattia Bosello, et al. thought that “heterobactin A” structure proposed by Carl J. Carrano, et al. is artificial due to technical issues. In the draft of latest SIDERITE, it’s planned to update “heterobactin A” to the structure proposed by Mattia Bosello, et al. However, we found another paper in 2021 in that another research group detected “heterobactin A” proposed by Carl J. Carrano, et al. in *R. erythropolis* S43 by LC-MS/HPLC^8^. Considering the strains used by Carl J. Carrano, et al. and Mattia Bosello, et al. are different and their “heterobactin A” co-occurring with different heterobactins, it’s possible that two “heterobactin A” structures both are correct. Therefore, in the latest SIDERITE, “heterobactin A” is split into “heterobactin A Carrano” proposed by Carl J. Carrano et al. and “heterobactin A Bosello” proposed by Mattia Bosello, et al.

### Figure S4

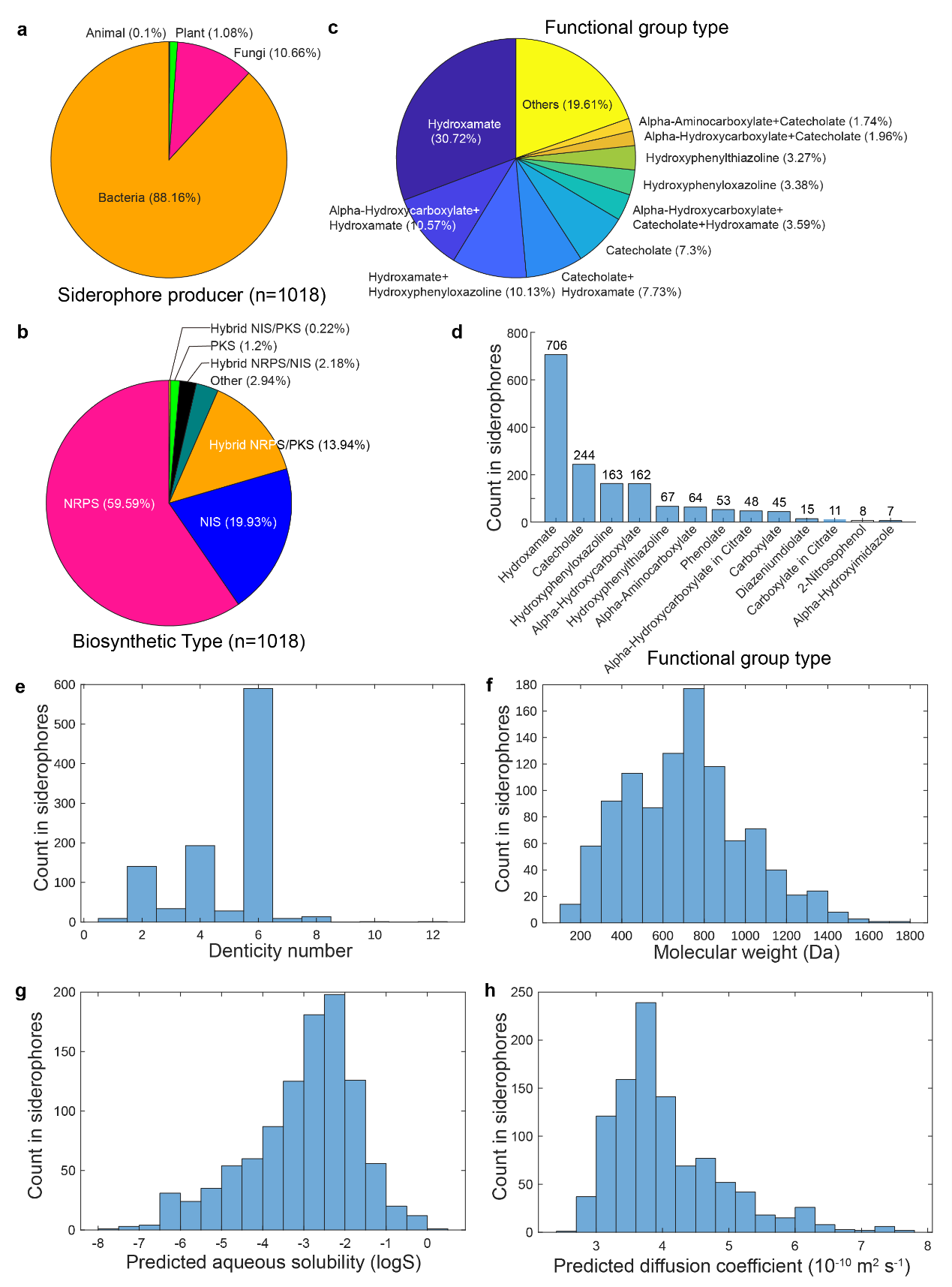

**Figure S4. The statistics of siderophore property.**

a. Distribution of the siderophore producers by their kingdoms.

b. Distribution of the siderophore biosynthetic pathways.

c. Distribution of the functional group type combinations. For clarity, only the top ten combinations are shown, and the others are merged into "Others".

d. Distribution of the common functional group of siderophores. One siderophore could contribute to more than one functional group type if it contains many types of functional groups.

e. Distribution of denticity numbers.

f. Distribution of the molecular weight.

g. Distribution of the predicted logS.

h. Distribution of the predicted diffusion coefficient (10^(-10) m2 s-1) in H2O, 298.15K.

### Figure S5

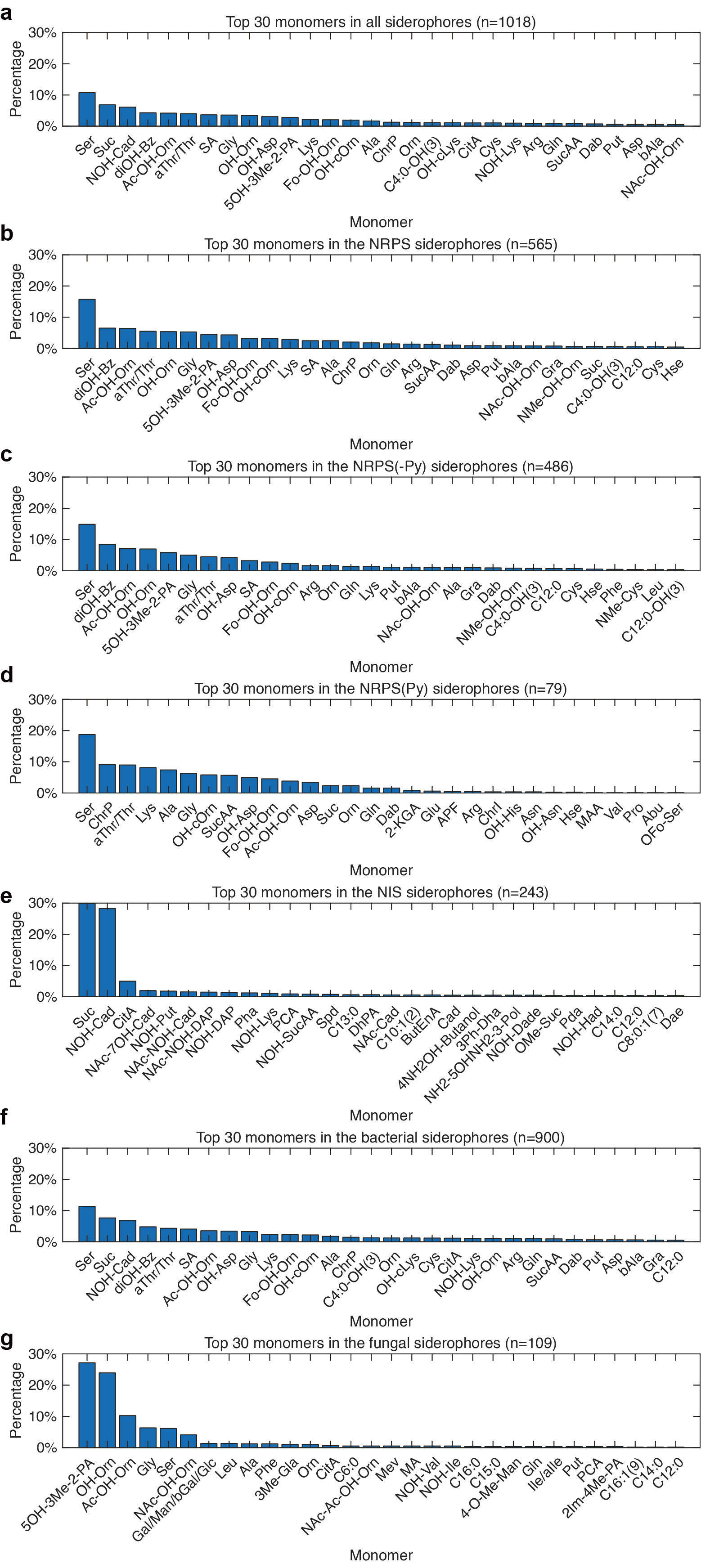

**Figure S5.** **The top 30 monomer distribution of siderophores.**

a. The top 30 monomer distribution of all siderophores.

b. The top 30 monomer distribution of NRPS siderophores.

c. The top 30 monomer distribution of NRPS siderophores without pyoverdines.

d. The top 30 monomer distribution of pyoverdine siderophores.

e. The top 30 monomer distribution of NIS siderophores.

f. The top 30 monomer distribution of bacterial siderophores.

g. The top 30 monomer distribution of fungal siderophores.
